# Assembly-free discovery of human novel sequences using long reads

**DOI:** 10.1101/2022.05.06.490971

**Authors:** Qiuhui Li, Bin Yan, Tak-Wah Lam, Ruibang Luo

**Affiliations:** Department of Computer Science, The University of Hong Kong, Hong Kong, China

**Author notes:** These authors contributed equally to this work (Qiuhui Li, Bin Yan). To whom correspondence should be addressed: R. L.

**Keywords:** Long reads, novel sequences, Assembly free approach, human references

## Abstract

DNA sequences that are absent in the human reference genome are classified as novel sequences. The discovery of these missed sequences is crucial for exploring the genomic diversity of populations and understanding the genetic basis of human diseases. However, various DNA lengths of reads generated from different sequencing technologies can significantly affect the results of novel sequences. In this work, we designed an Assembly-Free Novel Sequence (AF-NS) approach to identify novel sequences from Oxford Nanopore Technology long reads. Among the newly detected sequences using AF-NS, more than 95% were omitted from those using long-read assemblers, and 85% were not present in short reads of Illumina. We identified the common novel sequences among all the samples and revealed their association with the binding motifs of transcription factors. Regarding the placements of the novel sequences, we found about 70% enriched in repeat regions and generated 430 for one specific subpopulation that might be related to their evolution. Our study demonstrates the advance of the Assembly-Free approach to capture more novel sequences over other assembler based methods. Combining the long-read data with powerful analytical methods can be a robust way to improve the completeness of novel sequences.

## 1. Introduction

Building a complete reference genome in humans is fundamental for decoding genetic variation and the associations with human diseases ^1^. However, the current reference genome was derived from few individuals, resulting in an underrepresentation of the human population ^2,3^. To increase its diversity, human genome projects have assembled genomes across and within subpopulations ^4,5^. In general, DNA sequences missed in the human reference genomes are called novel sequences. These missed genomic sequences may contain thousands of unknown genetic variants implicated in biological functions. Therefore, identifying novel sequences can enrich the human reference genome and facilitate genome-based research.

Discovering novel sequences is dependent largely on the development of sequencing technologies. Initially, novel sequences were defined using fosmid end sequence pairs ^6^ and the entire fosmid clone ^7^. However, the high expense of capillary sequencing made the creation of large-scale genomes impractical. In contrast, Next-Generation Sequencing (NGS, also known as short-read sequencing) technologies enable the sequencing of thousands of samples with higher efficiency and at lower cost, resulting in more complete genomic information. Recently, an African pan-genome built with 910 African descents discovered 296 Mbp of novel sequences ^8^. We built a Chinese pan-genome using 486 Chinese genomes and got 276 Mbp of novel sequences ^9^. These studies have increased the human genome diversity and the related functional implications of novel sequences.

However, there is a major limitation for NGS short-reads <300 bases long that are too short to detect more variants, especially >70% of human genome structural variations ^10^. To overcome this shortcoming, long-read sequencing has emerged, including Oxford Nanopore Technology (ONT), which generates long (10-100 Kbp) and ultra-long (>100 Kbp) DNA reads ^11^. The average lengths of over 10,000 bp are helpful for analyzing structural variations and *de novo* assembly ^12-14^. The first gapless human reference was created using PacBio HiFi and ONT, which incorporated 200 Mbp sequences absent in GRCh38.p13 ^3^. Currently, detection of novel sequences relies mainly on genome assembling from long-read data. For example, 12.8 Mbp of novel sequences from a Chinese assembly ^15^ and over 10 Mbp of novel sequences from each individual in two Swedish genome projects ^16^. In 2019, a landmark work used 15 samples to produce the largest long-read structural variant callset and reported 6.4 Mbp of novel sequences per individual ^17^. These studies discovered a certain number of novel sequences, but considering varying lengths of DNA reads and different analytical approaches, it is necessary to evaluate their impact on building complete novel sequences.

In this study, we defined DNA sequences missed from the human reference and longer than 300 bp as “novel sequences”. We proposed an Assembly-Free Novel Sequence (AF-NS) approach that performs quick identification of novel sequences without assembling processes. Derived from ONT long-reads, the AF-NS detected novel sequences covered over 90% of Illumina novel sequences and contained more DNA information missing from the Illumina data. Our findings show the advantage of AF-NS in obtaining more intact DNA sequences while saving considerable computational resources, and the importance of long-read sequencing in building a complete picture of novel sequences. Furthermore, we uncovered the biological significance of the identified novel sequences and the population-specific novel insertions.

## 2. Materials and methods

### 2.1 Samples and data sources

To study the effects of sequencing technologies on detecting novel sequences, we selected both ONT long reads and Illumina paired-end short reads from six samples – an Ashkenazim trio (HG002, HG003, HG004) and a Chinese trio (HG005, HG006, HG007) – extracted from the Genome in a Bottle (GIAB) project ^18^. The ONT reads were base-called using Guppy v4.2.2. To keep the consistent sequencing depth for parent-offspring trios, we down-sampled the three Ashkenazim samples (HG002 to HG004) to 50-fold and the three Chinese samples (HG005 to HG007) to 40-fold. To guarantee the identified sequences/reads that are truly absent in the human reference, we combined the latest references, GRCh38.p13 and chm13.v1.1, as the new reference. In addition, we removed chromosome Y when identifying novel sequences of HG004 and HG007 because the two samples were female.

### 2.2 AF-NS: assembly-free novel sequences construction

As shown in Figure 1, AF-NS carried out an assembly-free pipeline consisting of three main steps. First, we aimed to obtain high-quality unmapped reads. All ONT long reads were aligned to references using minimap 2.17-r941^19^, and reads with unmapped fragments longer than 300bp were selected. We then discarded low-quality reads with <10 Q-score by NanoFilt ^20^, and cut the adaptors of reads using porechop v0.2.4 ^21^. Second, we collected the unmapped sequences by three-round alignment-filtering. The filtered reads were aligned to references using minimap2. The unmapped fragments were then aligned to references using NUCmer 4.0.0rc1^22^. The mapped fragments were removed, and the rest were aligned to references, again using minimap2. Any fragments with alignments to references and shorter than 300bp were removed. Third, we eliminated sequences labelled as archaea, bacteria, fungi, plasmid, viral and UniVec (confidence score >0.05) by Kraken2 ^23^ and used minimap2 to overlap the remaining sequences. Two sequences were clustered together if they aligned with each another with >80% coverage. We chose the longest sequence in a cluster as its representative. Since the long-read sequencing could be inaccurate in low-complexity regions ^24^, we utilized RepeatMasker 4.1.2-p1 ^25^ to annotate the novel sequences and removed sequences if 80% of them were low-complexity or simple repeats. The remaining ones were collected as the final set of novel sequences.

**Figure 1.**
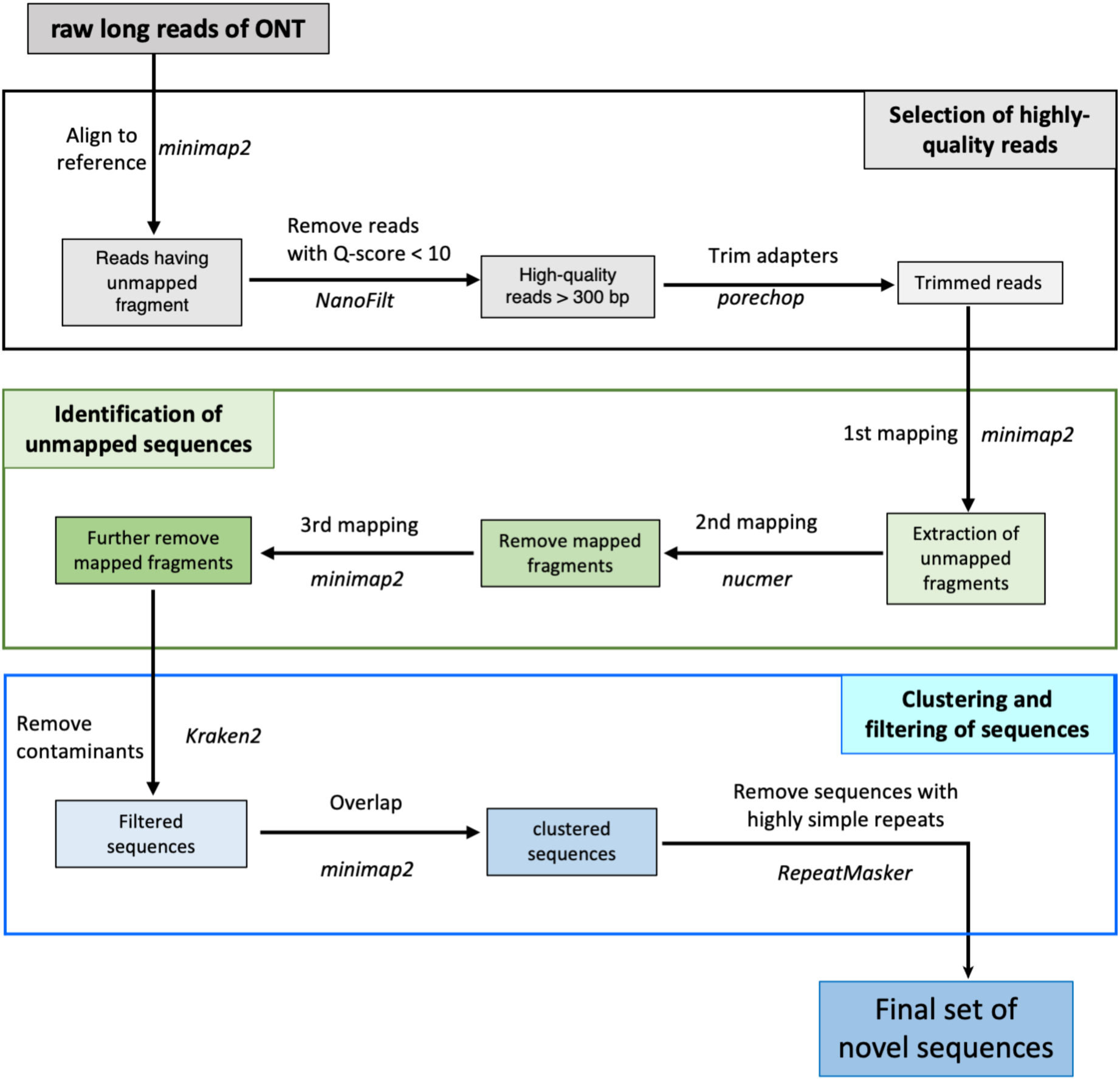
Workflow for identification of long-read novel sequences by Assemble-free Novel sequence (AF-NS) approach. Using long-reads of Oxford Nanopore Technology (ONT), AF-NS carries out three main steps

### 2.3 novel_WG: extraction of novel sequences from the whole-genome assembly

We executed whole genome assemblies using two popular assemblers, Shasta 0.7.0 ^26^ and raven 1.7.0 ^27^, and aligned the assemblies to references using NUCmer. Any fragments mapped to the reference with ≥ 80% identity were removed. Next, we aligned the unmapped fragments >300 bp to references using BWA-mem v0.7.17 ^28^ and further eliminated the mapped parts if the alignment identity reached 80%. Then, to ensure the novelty of the remaining sequences, we aligned them to references using BWA-mem again and discarded any sequences that had hits to references satisfying >80% identity and >50% coverage. Sequentially, we used Kraken2 to remove contaminated sequences to obtain the final set of novel sequences. For Illumina data, we performed *de novo* assembly using MEGAHIT v1.2.9 ^29^. Other procedures were the same as those used for long-read sequencing data.

### 2.4 unmap_ASM: assembly of unmapped reads

Illumina reads were mapped to human references using BWA-mem, and the unmapped parts were obtained and assembled into sequences using MEGAHIT. Next, we deleted novel sequences shorter than 300 bp and aligned the remaining ones to references using BWA-mem. Any sequences mapped to references with identity ≥ 80% and coverage ≥ 50% were filtered out. Finally, we eliminated contaminants identified by Kraken2 and generated a set of novel sequences.

### 2.5 Detection of novel placements

We aligned the ONT long reads to chm13 v1.1 using minimap2 and retained alignments with ≥ 80% identity. For reads with 1 or 2 consistent alignments, we extracted the corresponding unmapped fragments longer than 1000 bp. These fragments were classified into two groups: (1) those with a single end aligned to the reference (SEP), and (2) those with both ends mapped to the reference (BEP). Then we clustered two types of fragments. Two BEP sequences were merged if the length difference between their overlap and length was less than 100 bp. SEP sequences within 100 bp of each other were combined using BEDtools ^30^. The longest sequence in a cluster was selected as its representative. We excluded clusters containing fewer than three sequences to obtain the final placement clusters.

## 3. Results

### 3.1 Procedure for detecting long-read novel sequences

To examine the effects of different length reads on the discovery of novel sequences, we chose six samples, an Ashkenazim trio (HG002, HG003, HG004) and a Chinese trio (HG005, HG006, HG007), both of which have ONT long-read and Illumina short-read data. First, we conducted two widely used strategies to acquire long-read novel sequences, named novel_WG and unmap_ASM. The first approach involved extracting novel sequences from the whole-genome assembly. We used Shasta and raven to generate the whole-genome assemblies. However, the resulting novel sequences differed greatly in size, and more than 50% of them could not be aligned against each other with ≥ 80% identity (Table1, Supplementary Table 1). This inconsistency might be due to different whole-genome assemblers. The authenticity of these unmapped sequences requires further verification to determine whether there are assembly-caused artifacts. The mapped ones were treated as common novel_WG sequences with high confidence. Using the unmap_ASM approach, we obtained only a few novel sequences, indicating its ineffectiveness for long-read data (Supplementary Table 2).

To discover more complete long-read novel sequences, we designed a new pipeline AF-NS, consisting of a three-step procedure without using assemblers (Figure 1). The first step obtained high-quality reads unmapped to the human reference and trimmed their adaptors. Then, through a three-round alignment-filtering process, the unmapped fragments longer than 300 bp were extracted from the trimmed reads as novel candidates. During the last step, after removing contaminants, similar candidates were clustered together to generate the final set of novel sequences. Compared with other methods, which assemble reads into contigs (contiguous sequences), AF-NS obtains novel genomic information from high-quality long reads. This ensures its efficiency in detecting novel sequences through minimizing sequence loss due to assembling.

### 3.2 Comparison of novel sequences

First, we compared AF-NS with the assembly-based approach novel_WG. AF-NS does not require computationally intensive whole genome assembly, and can detect novel sequences 15 times more than novel_WG (Table 1). We note that over 85% of the common novel_WG sequences were mapped to AF-NS identified sequences with ≥ 80% identity (Table 2), showing the superiority of AF-NS in capturing more novel genomic information. Then we aligned the AF-NS sequences to whole-genome assemblies using NUCmer. The mapping percentage was less than 5% with ≥ 50% coverage (Table 3). This result indicates that current long-read assemblers lose most of the novel genomic information.

**Table 1.** Size (bp) of long-read novel sequences identified by AF-NS and novel_WG.

| Samples | AF-NS | novel_WG (Shasta) | novel_WG (raven) |
| --- | --- | --- | --- |
| HG002 | 19,055,769 | 182,032 | 1,328,545 |
| HG003 | 23,950,540 | 207,148 | 728,462 |
| HG004 | 22,731,658 | 240,347 | 2,141,062 |
| HG005 | 15,973,628 | 224,995 | 1,361,765 |
| HG006 | 16,549,640 | 167,900 | 1,201,172 |
| HG007 | 15,853,142 | 238,687 | 1,024,175 |

AF-NS is a newly designed Assembly-free approach. novel\_WG detects novel sequences from the long-read whole genome assembly generated using two assemblers, Shasta and raven, respectively.

**Table 2.** Comparison of long-read novel sequences (bp) using AF-NS and novel_WG.

| Samples | common novel_WG vs. AF-NS sequences<br>aligned length (percentage of alignment) | AF-NS sequences vs. common novel_WG<br>aligned length (percentage of alignment) |
| --- | --- | --- |
| HG002 | 59,429 (86.56%) | 480,109 (2.52%) |
| HG003 | 58,160 (95.7%) | 416,477 (1.74%) |
| HG004 | 91,127 (93.85%) | 769,500 (3.39%) |
| HG005 | 73,791 (85.76%) | 360,576 (2.26%) |
| HG006 | 73,261 (91.51%) | 405,168 (2.45%) |
| HG007 | 100,790 (90.99%) | 557,672 (3.52%) |

Common novel\_WG: the novel sequences extracted from both Shasta and raven assemblies using novel\_WG approach can be aligned with each other with $\geq 80\%$ identity. AF-NS sequences: long-read novel sequences identified by the AF-NS approach.

**Table 3.**
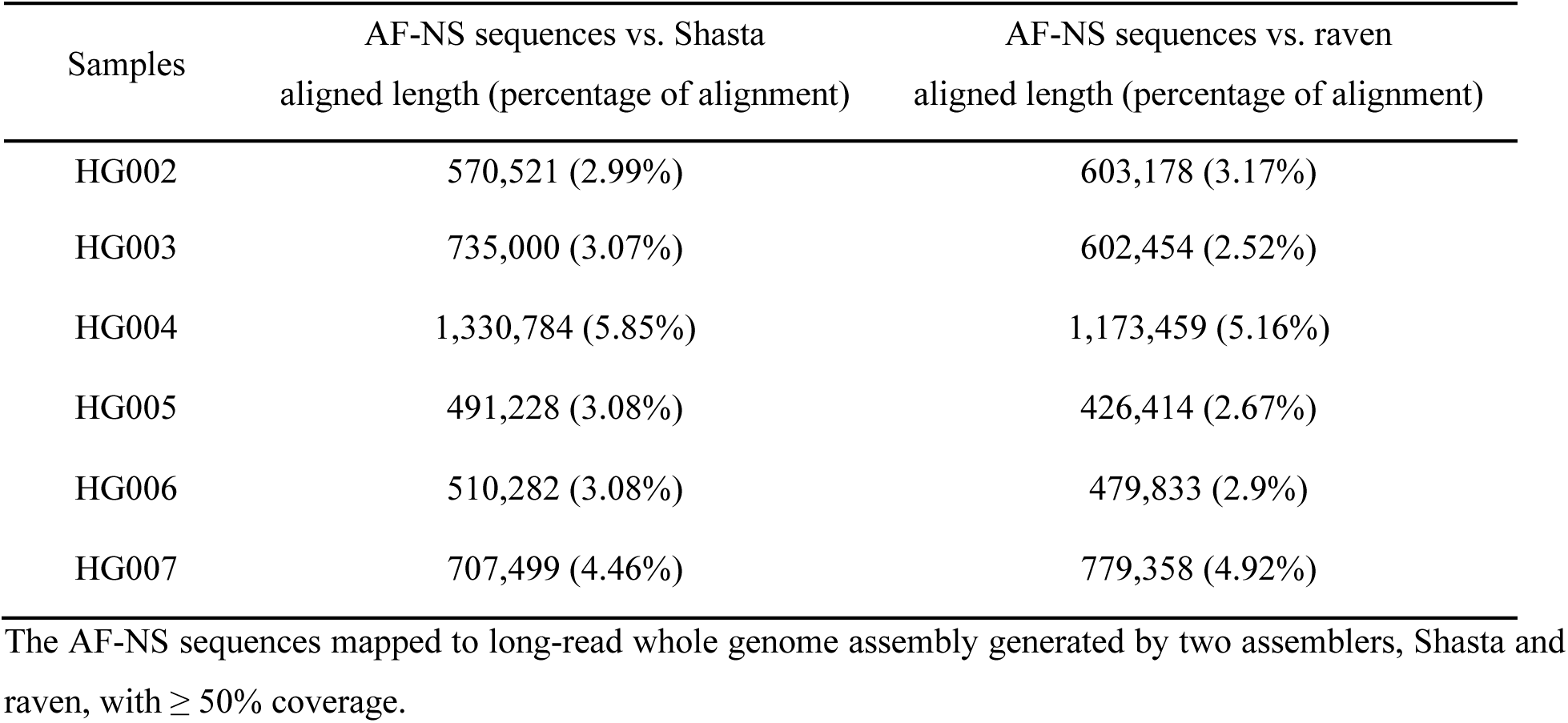
Alignment of AF-NS sequences to long-read whole genome assembly (bp)

| Samples | AF-NS sequences vs. Shasta | AF-NS sequences vs. raven |
| --- | --- | --- |
|  | aligned length (percentage of alignment) | aligned length (percentage of alignment) |
| HG002 | 570,521 (2.99%) | 603,178 (3.17%) |
| HG003 | 735,000 (3.07%) | 602,454 (2.52%) |
| HG004 | 1,330,784 (5.85%) | 1,173,459 (5.16%) |
| HG005 | 491,228 (3.08%) | 426,414 (2.67%) |
| HG006 | 510,282 (3.08%) | 479,833 (2.9%) |
| HG007 | 707,499 (4.46%) | 779,358 (4.92%) |

The AF-NS sequences mapped to long-read whole genome assembly generated by two assemblers, Shasta and raven, with $\geq 50\%$ coverage.

Next, we compared the novel sequences generated from different sequencers. We first constructed Illumina novel sequences of the six samples using two methods, unmap_ASM and novel_WG. The size of novel_WG sequences was slightly larger than that of unmap_ASM sequences (Table 4). We aligned novel_WG and unmap_ASM sequences with each other and found that ∼ 85% of unmap_ASM sequences or ∼ 95% of novel_WG sequences could be mapped together with ≥ 80% identity (Supplementary Table 3). This result shows the consistency of novel sequences obtained from the two approaches. To minimize assembly-induced artifacts, we retained only the novel sequences detected by both. The HG005 sample was excluded because the size of novel sequences discovered from its Illumina reads was considerably higher than that of the other samples (Table 4).

**Table 4.**
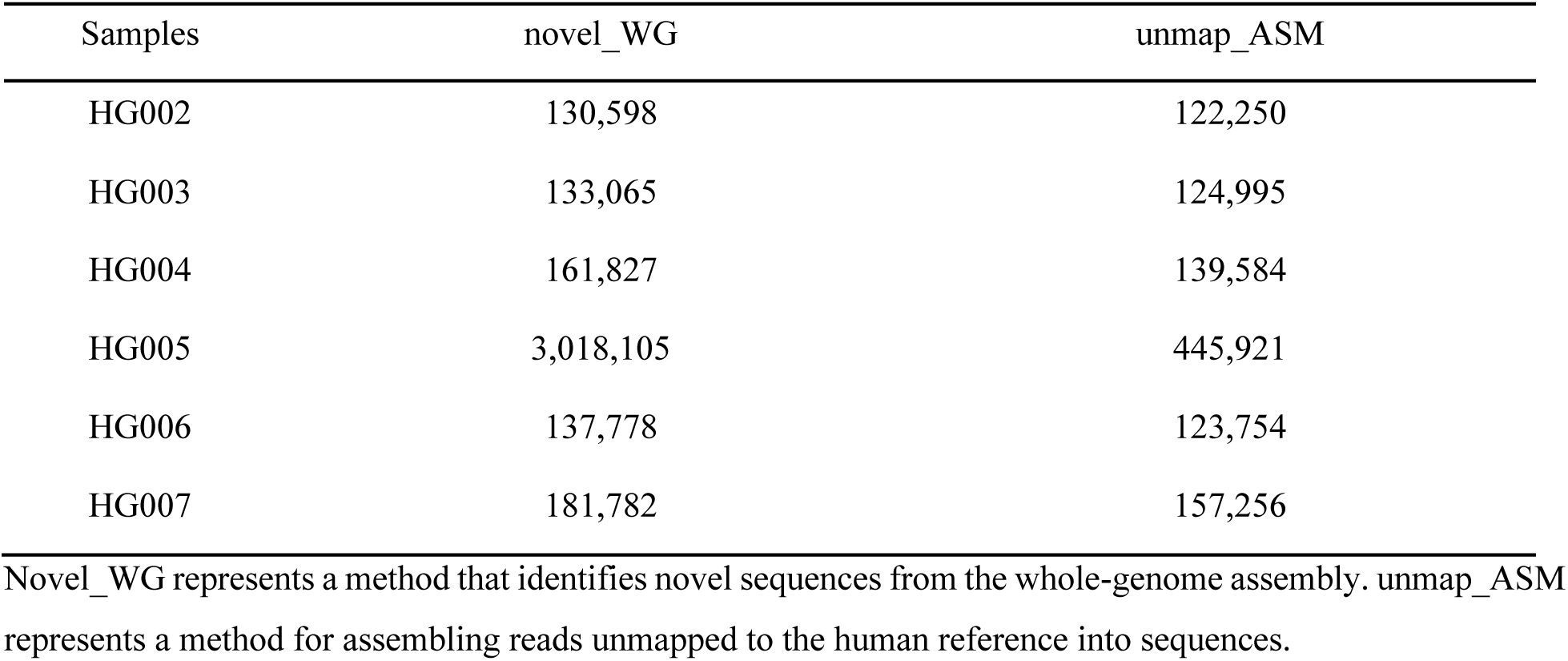
Size (bp) of short-read novel sequences identified by two methods.

As shown in Figure 2A, the AF-NS identified sequences covered more than 91% of Illumina novel sequences under ≥ 80% identity and ≥ 50bp alignment length. By contrast, only 58-80% of the Illumina novel sequences were mapped to the long-read whole genomes using an assembler Shasta or raven (Figure 2A). This analysis verifies the effectiveness of the AF-NS approach in discovering intact novel sequences. Because the above two identification methods may lose some novel sequences, we mapped entire Illumina raw reads to the AF-NS sequences using BWA-mem. Under the same threshold, less than 15% of the AF-NS sequences were covered by Illumina data (Figure 2B). We also ran BUSCO ^31^ on both ONT and Illumina whole-genome assemblies using the eukaryote set of orthologs from OrthoDB v.10 ^32^. As expected, the short-read whole genome was not as complete as the long-read assemblies (Supplementary Table 4). These results indicate that short-read sequencing does miss a lot of genomic information.

**Figure 2.**
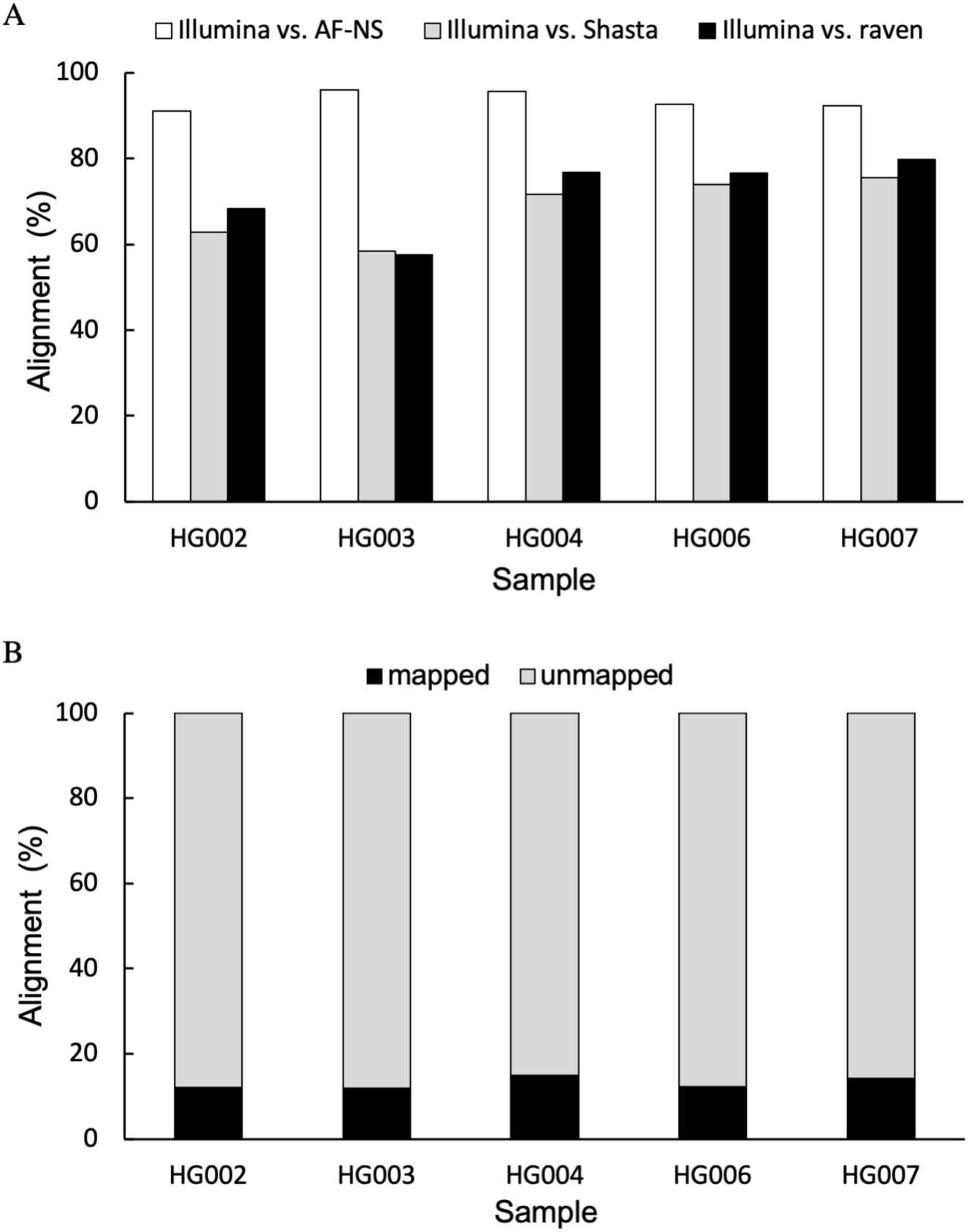
Comparison of short-read and long-read novel sequences. A, the percentage of short-read novel sequences of Illumina that were aligned to long-read sequences; AS-NS: long-read novel sequences identified by the AS-NS method; Shasta: long-read whole-genome assemblies generated using Shasta; raven: long-read whole-genome assemblies generated using Shasta. B, the percentage of long-read novel sequences that were mapped to short-read raw reads; mapped: long-read novel sequences were mapped to short-read raw reads; unmapped: long-read novel sequences were not mapped to short-read raw reads. All alignments were based on

### 3.3 Characteristics of novel sequences

According to our previous definition of the novel sequence composition ^9^, we classified the AF-NS detected sequences as common and individual-specific components. A sequence was considered common if it could be mapped to other samples’ sequences by satisfying ≥ 80% identity. Each of the six samples contained 1-2 Mbp common sequences, accounting for 6-8% of total sequences (Supplementary Table 5). To search for the origin of the AF-NS sequences, we aligned them to the chimpanzee genome (GCF_002880755.1) using NUCmer. Under the minimum requirement of 80% identity, about 7% of the novel sequences were present in the chimpanzee genome. By contrast, the common sequences of each sample obtained much higher alignment percentages of ∼ 60% compared to individual-specific sequences of 2-3% (Figure 3A).

**Figure 3.**
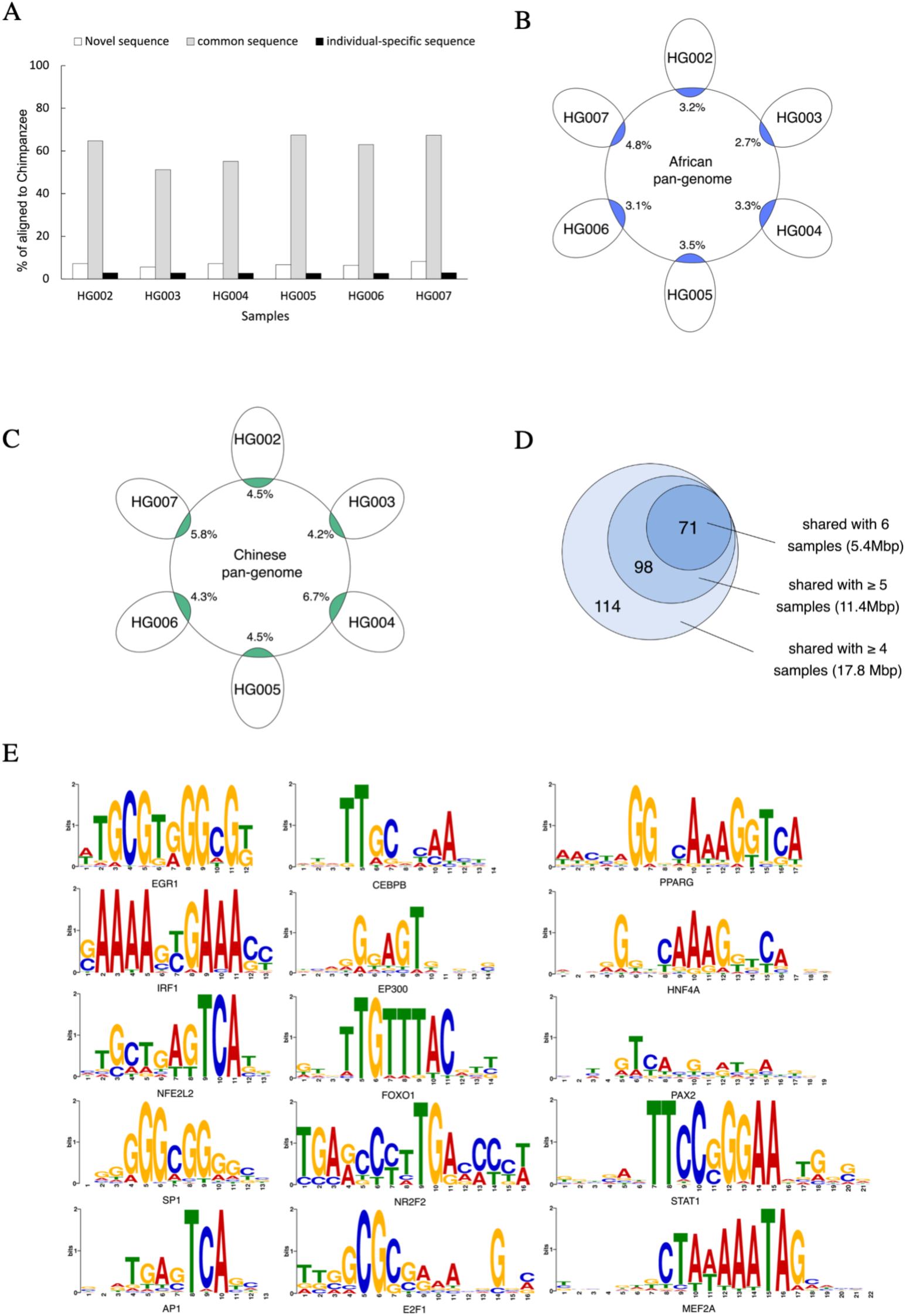
Analysis of the characteristics of long-read novel sequences. A, percentage of the AF-NS identified novel sequences mapped to the chimpanzee genome with ≥ 80% identity. Common sequence: novel sequences shared by at least two samples. Individual-specific sequence: novel sequences found in one individual sample. B-C, percentage of the AF-NS identified sequences mapped to African pan-genome (B) and Chinese pan-genome (C) with ≥ 80% identity. D, number of transcription factor (TF) binding motifs identified from the highly common sequences that are shared among at least four, five and six samples, respectively. The lengths of highly common sequences are also shown. E, binding motif logos of 15 main TFs that were predicted to bind to the

To determine the novelty of the AF-NS sequences, we mapped them to existing novel sequences of three long-read sequencing samples, HX1 (Chinese) and two Swedes ^15,16^. Few AF-NS sequences were aligned with ≥ 80% identity, indicating that most of them have not been reported (Supplementary Table 6). Next, we compared the AF-NS sequences with two pan-genomes, the African pan-genome and the Chinese pan-genome ^33,34^. Only about 3-7% of the AF-NS sequences could be matched to either one, implying a great number of novel sequences lost in the pan-genomes (Figures 3B and 3C). The low alignment rate is possibly because the current pan-genomes were assembled using short-read data. However, the percentage of alignment to pan-genomes was still 3-4 times higher than that to the long-read novel sequences of an individual, emphasizing the importance of large-scale sequencing and assembling for obtaining complete genomic information.

In order to reveal the biological significance of the identified novel sequences, we respectively extracted the highly common sequences shared among at least four, five and all six samples. Transcription factors (TFs) represent the main regulators that control gene transcription by binding to the DNA sequences. We performed TF binding motif searching on the highly-common sequences using fimo ^35^ and based it on the TRANSFAC human dataset ^36^. With a cutoff of p-value < 5e-8, we searched for TF binding to these highly common sequences. There were 71 TF motifs identified to bind to the sequences shared by all six samples (Figure 3D, Supplementary Table 7). We display binding patterns of 15 main TFs that are believed to have a broad set of biological functions (Figure 3E), such as transcriptional regulation (CEBPB, P300), immune or inflammatory response (AP1, IRF1, STAT1, PPARG, NFE2L2), tumors (AP1, SP1, EGR1, STAT1, PAX2, etc.), metabolism (FOXO1, PPARG), and growth and development of cells or organs (E2F1, SP1, MEF2A, NR2F2, HNF4A, PAX2, NEF2L2, PPARG, etc.). These proposed TFs suggest possible functional implications of our novel sequences.

### 3.4 Analysis of novel placements

We positioned novel sequences in chm13.v1.1 and allowed the placed sequences within 100 bp of each other to be clustered together. We obtained 1,637 placed clusters among the six samples and used the corresponding placements to analyze the inserting preferences. The majority of placements (69%) were located in repeat regions, especially satellite and microsatellite regions (Supplementary Table 8, Figure 4A). The enrichment of novel sequences in repeats might be due to the highly unstable nature of these regions, which are prone to generating variants. These variations probably drive genome plasticity and promote evolution ^37,38^. In addition, we found 54 and 68 novel placements were in centromeres and telomeres, respectively, where genomic information is difficult to obtain using short-read data ^39^. Long-read sequencing enables the mapping of novel sequences in highly condensed heterochromatin regions.

**Figure 4.**
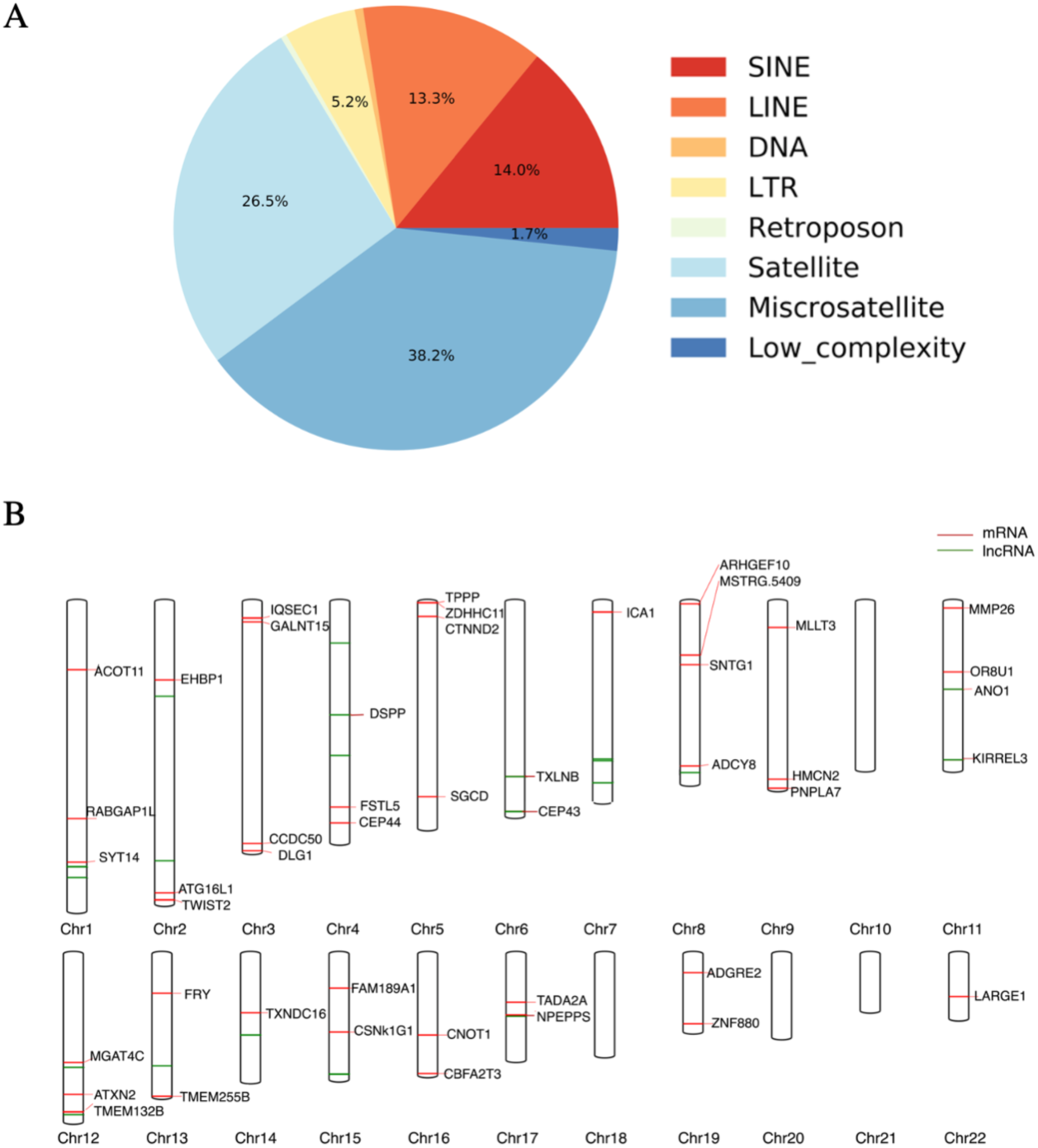
Properties of novel placements. A, percentage of novel sequence placements located in interspersed repeat and low-complexity regions. Satellites and microsatellites account for the largest proportions. B, in the genome map, the red and green lines represent the novel sequence placements located in protein-coding and lncRNA regions on autosomes, respectively. These placements are shared among all samples. The corresponding protein-coding gene symbols are shown. Detailed annotations are reported in Supplementary Table 8.

Among the novel placements, 1,514 common ones were present in at least two samples (Figure 5A). Since the GIAB samples were from Ashkenazim and Chinese subpopulations, we further divided the common novel placements into population-shared (across subpopulations) and population-specific (within a subpopulation). The 1,084 population-shared insertions as patches can supplement the latest reference genome. We obtained 430 population-specific insertions, including 198 Chinese-specific and 232 Ashkenazim-specific (Figure 5B) insertions. Then, we found 260 population-specific insertions present in only one parent and its offspring. If these mutations are newly generated in the parents, they may serve as new heritable variants.

**Figure 5.**
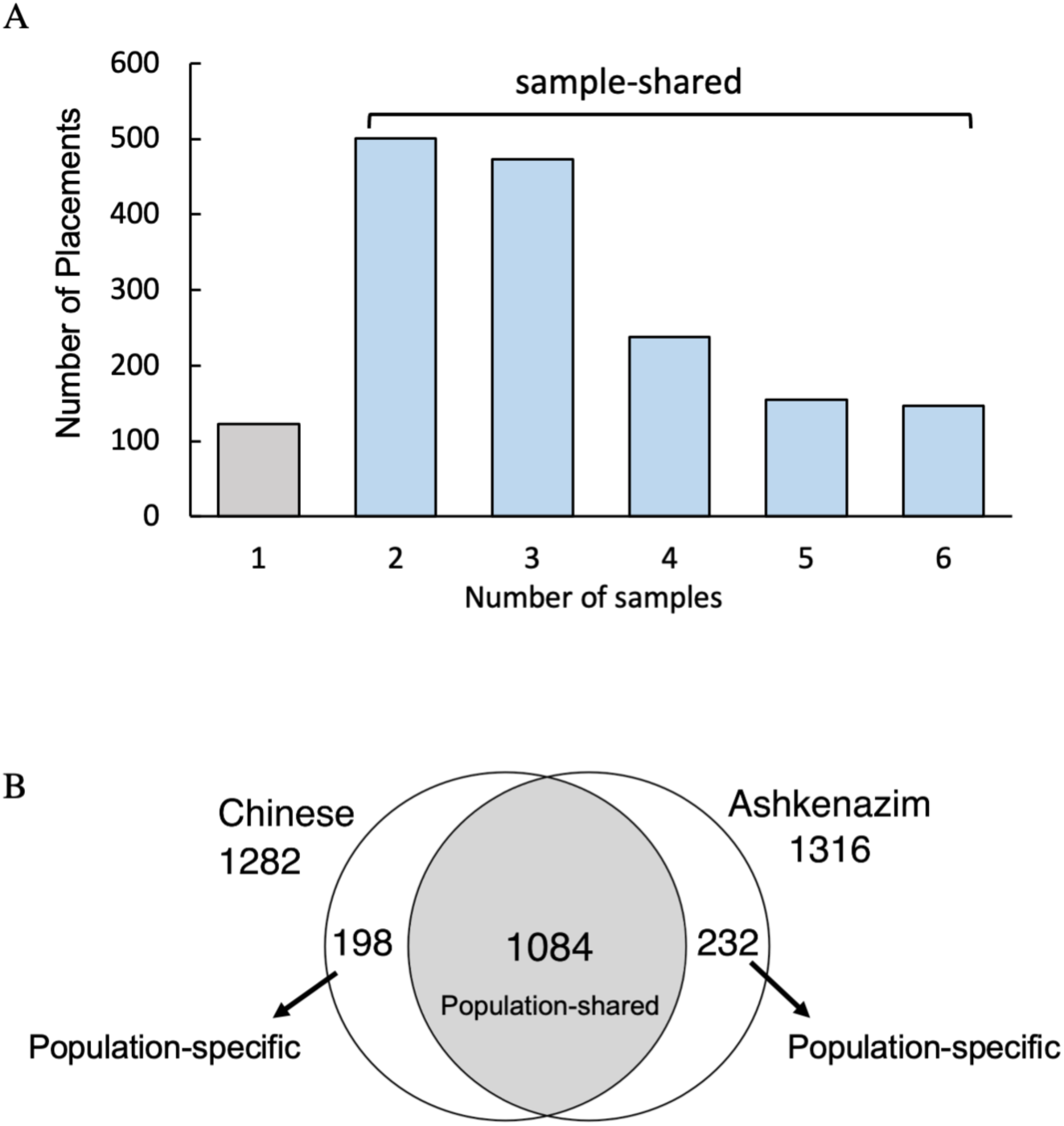
Novel sequence placements detected in six samples. A, number of novel sequence placements detected in only one or shared by two to six samples. B, intersection of the novel sequence placements shared by at least two samples within the Ashkenazim or Chinese subpopulation.

Furthermore, we investigated the functional significance of the placed insertions using chm13 gene annotation (version4). A total of 613 insertions fell within 503 genes, including 301 coding and 162 long non-coding (lncRNA) regions (Supplementary Table 8). We reported 46 protein-coding and 22 lncRNA genes shared among all samples (Figure 4B). The 46 protein-coding genes involved mainly cell adhesion (*ADGRE2, CTNND2, HMCN2, DSPP, KIRREL3* and *DLG*), actin cytoskeleton organization (*ARHGEF10, IQSEC1* and *EHBP1*), endocytosis (*RABGAP1L, EHBP1* and *CSNK1G1*), and multicellular organism development (*ANO1, FSTL5, TWIST2* and *DSPP*).

## 4. Discussion

The identification of novel sequences is a basis for complementing the existing reference genome in humans ^40^. However, influenced by different sequencing technologies and computational methods, the derived novel sequences vary considerably ^16,41,42^. Finding a widely accepted standard for constructing novel sequences is a challenging issue in genomics research. Currently, there are two strategies for building novel sequences. The first one involves filtering out reads mapped to references and assembling the unmapped reads into novel sequence sets ^8,9,43^. Another involves performing the whole-genome *de novo* assembly and extracting the unmapped sequences ^16,42^. However, the quality of the identified novel sequences using these strategies is highly dependent on the performance of assemblers. When assembling unmapped reads, some novel sequences can be lost because of insufficient supporting reads or overlaps between reads that fail to meet assembly requirements^44^. Owing to the short length of NGS data, genome assembly is essential for identifying novel sequences that exceed the read lengths. By contrast, long-read sequencing can generate reads longer than 100 Kbp. Therefore, a single read can decipher large structural variations, which guarantees the feasibility of AF-NS to discover novel sequences at read level. Our assembly-free strategy includes a series of processes to ensure the completeness and accuracy of the identified novel sequences. First, the selected ONT reads during the first step should be of high quality. Second, the contaminants in the clustered candidate sequences must be filtered out. Last, we removed highly simple repetitive sequences from the final set due to the high error rate of long-read sequencing in low-complexity regions ^24^. The performance of AF-NS demonstrates its advances in identifying novel sequences that have a larger size and better characteristics over other methods, and saves considerable computational resources.

With the advancement of sequencing technologies, different ones have been applied to detect novel sequences ^15,41,45^. However, the corresponding analyses focused only on specific datasets. Here, we compared the novel sequences generated using short- and long-read data and found a big difference, i.e., that most of the long-read sequences cannot be covered by short-read ones. Also, most of the long-read novel sequences were absent in Illumina pan-genomes even though the pan-genomes were built using hundreds of samples. These findings demonstrate that utilizing long-read technologies is the inevitable trend for creating complete novel DNA sequences.

Interaction between TFs and TF binding DNA sites on promotors is a key for transcriptional gene regulation. To analyze biological associations of the AF-NS novel sequences, we identified 71 TF binding motifs on the highly common sequences shared by all samples. These TFs are assumed to play regulatory functions involving many basic biological processes and pathways. Although the TF binding sites were not confirmed to promoters or noncoding regions, this finding suggests a wide biological connection to the newly detected sequences. Moreover, we discovered the population-shared and population-specific insertions by clustering placed novel sequences of six samples. The population-shared insertions indicate that the latest human reference genome is still underestimated, possibly because chm13 was derived from one genome ^3^. The population-specific insertions may be the force of the subpopulation evolution. Considering our classification using only six samples, increasing individuals is expected to assist in establishing more representative population-specific or common insertions. These population-specific insertions will probably allow us to determine the relationships of novel sequences with human evolution.

In summary, AF-NS represents a rapid sequence detection method from long reads and outperforms the existing assemblers in discovering intact novel sequences. The assembly-free design considerably reduces computation costs and facilitates the construction of large-scale novel sequence genomes. The superiority of long-read sequencing in identifying more novel sequences and positioning them in difficult-to-assemble regions, like centromeres and telomeres, shows its great potential for future novel sequence research. The high cost of long-read sequencing is still a challenge for generating numerous genomes. However, as more samples using advanced sequencers become available, we believe that we can establish a more comprehensive landscape of novel sequences.

## Supplementary data

SupplementaryData.zip

## Data availability

The AF-NS identified sequences can be found at https://www.bio8.cs.hku.hk/novel/AF-NS/.The implementation of the AF-NS method is available at https://github.com/HKU-BAL/AF-NS.

The Illumina paired-end short reads of HG002-HG007 were downloaded from https://github.com/genome-in-a-bottle/giab_data_indexes. The ONT long reads of HG002-HG007 were downloaded from https://s3-us-west-2.amazonaws.com/human-pangenomics/index.html?prefix=NHGRI_UCSC_panel/. The novel sequences of 910 Africans in APG were downloaded from NCBI with accession number PDBU01000000. The novel sequences of 486 Chinese in CPG were downloaded from the Genome Sequence Archive for Human with the accession number PRJCA003657. The assembly of HX1 was downloaded from http://www.openbioinformatics.org/hx1/data/nonhg38.fa.gz. The assemblies of the two Swedes were downloaded from https://www.mdpi.com/2073-4425/9/10/486 in Supplementary Data S1.

## Acknowledgement

R.L. was supported by the General Research Funding [17113721] of the Hong Kong SAR government, General Program [JCYJ20210324134405015] of the Shenzhen municipal government, China, and URC fund by the University of Hong Kong.

## Authors’ contributions

Q. L., and B. Y. performed data processing and analysis. R. L. conceived and advised the project. All authors wrote and approved the final manuscript.

