## Supplementary Materials for "Assembly-free discovery of human novel sequences using long reads"

**Supplementary Table 1. Alignment of the novel_WG identified novel sequences using two assemblers Shasta and raven (bp)**

| Samples | Shasta vs. raven<br>aligned length (percentage of alignment) | raven vs. Shasta<br>aligned length (percentage of alignment) |
|---|---|---|
| HG002 | 68,653 (37.71%) | 66,099 (4.98%) |
| HG003 | 60,776 (29.34%) | 172,904 (23.74%) |
| HG004 | 97,102 (40.4%) | 98,200 (4.59%) |
| HG005 | 86,047 (38.24%) | 159,546 (11.72%) |
| HG006 | 80,059 (47.68%) | 79,039 (6.58%) |
| HG007 | 110,768 (46.41%) | 110,493 (10.79%) |

Alignment of the novel sequences using Shasta and raven must meet at least 80% identity.

**Supplementary Table 2. Size (bp) of unmapped_ASM identified long-read novel sequences using two assemblers Shasta and raven**

| Samples | novel sequences<br>assembled by raven | novel sequences<br>assembled by Shasta |
|---|---|---|
| HG002 | 168,804 | 0 |
| HG003 | 360,027 | 0 |
| HG004 | 317,681 | 6,189 |
| HG005 | 209,750 | 0 |
| HG006 | 834,088 | 0 |
| HG007 | 262,603 | 0 |

**Supplementary Table 3. Alignment of the short-read novel sequences identified by novel_WG and unmap_ASM (bp)**

| Samples | novel_WG vs. unmap_ASM<br>aligned length (percentage of alignment) | unmap_ASM vs. novel_WG<br>aligned length (percentage of alignment) |
|---|---|---|
| HG002 | 117,898 (90.28%) | 116,307 (95.14%) |
| HG003 | 119,188 (89.57%) | 118,456 (94.77%) |
| HG004 | 136,944 (84.62%) | 133081 (95.34%) |
| HG006 | 119,585 (86.8%) | 116,689 (94.29%) |
| HG007 | 153,800 (84.61%) | 151,222 (96.16%) |

Alignment of the novel sequences must meet at least 80% identity

**Supplementary Table 4. BUSCO results of three whole genome assemblies**

| Sample | Metric | megahit | raven | Shasta |
|---|---|---|---|---|
| HG002 | Complete BUSCOs (C) | 76.40% | 98.10% | 97.60% |
| | Complete and single-copy BUSCOs (S) | 67.80% | 91.40% | 93.30% |
| | Complete and duplicated BUSCOs (D) | 8.60% | 6.70% | 4.30% |
| | Fragmented BUSCOs (F) | 22.40% | 1.60% | 2.00% |
| | Missing BUSCOs (M) | 1.20% | 0.30% | 0.40% |
| HG003 | Complete BUSCOs (C) | 75.60% | 98.80% | 98.40% |
| | Complete and single-copy BUSCOs (S) | 67.80% | 89.40% | 92.50% |
| | Complete and duplicated BUSCOs (D) | 7.80% | 9.40% | 5.90% |
| | Fragmented BUSCOs (F) | 21.60% | 0.80% | 1.60% |
| | Missing BUSCOs (M) | 2.80% | 0.40% | 0.00% |
| HG004 | Complete BUSCOs (C) | 75.70% | 98.80% | 98.40% |
| | Complete and single-copy BUSCOs (S) | 67.50% | 92.50% | 89.80% |
| | Complete and duplicated BUSCOs (D) | 8.20% | 6.30% | 8.60% |
| | Fragmented BUSCOs (F) | 20.80% | 1.20% | 1.60% |
| | Missing BUSCOs (M) | 3.50% | 0.00% | 0.00% |
| HG005 | Complete BUSCOs (C) | 54.90% | 98.50% | 98.10% |
| | Complete and single-copy BUSCOs (S) | 48.60% | 91.80% | 91.00% |
| | Complete and duplicated BUSCOs (D) | 6.30% | 6.70% | 7.10% |
| | Fragmented BUSCOs (F) | 36.90% | 0.80% | 2.00% |
| | Missing BUSCOs (M) | 8.20% | 0.70% | 0.10% |
| HG006 | Complete BUSCOs (C) | 75.30% | 99.30% | 98.50% |
| | Complete and single-copy BUSCOs (S) | 67.10% | 91.80% | 92.20% |
| | Complete and duplicated BUSCOs (D) | 8.20% | 7.50% | 6.30% |
| | Fragmented BUSCOs (F) | 22.70% | 0.40% | 1.20% |
| | Missing BUSCOs (M) | 2.00% | 0.30% | 0.30% |
| HG007 | Complete BUSCOs (C) | 73.80% | 97.60% | 98.90% |
| | Complete and single-copy BUSCOs (S) | 66.30% | 89.80% | 92.20% |
| | Complete and duplicated BUSCOs (D) | 7.50% | 7.80% | 6.70% |
| | Fragmented BUSCOs (F) | 24.70% | 2.00% | 1.20% |
| | Missing BUSCOs (M) | 1.50% | 0.40% | 0.10% |

The assemblies are compared with the eukaryote set of orthologs. Short-read whole genome was assembled by megahit. Long-read whole genome assembled by raven and Shasta, respectively.

**Supplementary Table 5. Size (bp) of long-read common sequences identified by AF-NS**

| Samples | Common sequence length (percentage of common sequences) |
|---|---|

| | |
|---|---|
| HG002 | 1,351,058 (7.09%) |
| HG003 | 1,392,666 (5.81%) |
| HG004 | 1,947,748 (8.57%) |
| HG005 | 1,009,966 (6.32%) |
| HG006 | 1,030,524 (6.23%) |
| HG007 | 1,308,942 (8.26%) |

Common sequences: the novel sequences shared by at least two samples based on minimal 80% identity.

**Supplementary Table 6. Size (bp) and percentage of the AF-NS identified long-read novel sequences aligned to the three individuals**

| Samples | Nanopore vs. HX1 | Nanopore vs. Swe1 | Nanopore vs. Swe2 |
|---|---|---|---|
| HG002 | 107,789 (0.57%) | 230,362 (1.21%) | 194,758 (1.02%) |
| HG003 | 141,152 (0.59%) | 205,983 (0.86%) | 236,210 (0.99%) |
| HG004 | 131,668 (0.58%) | 240,413 (1.06%) | 201,234 (0.89%) |
| HG005 | 105,655 (0.66%) | 159,782 (1.0%) | 170,975 (1.07%) |
| HG006 | 105,320 (0.64%) | 180,085 (1.09%) | 184,491 (1.11%) |
| HG007 | 111,931 (0.71%) | 186,465 (1.18%) | 224,455 (1.42%) |

Alignment of the novel sequences must meet at least 80% identity

**Supplementary Table 7. A list of transcription factor (TF) binding motifs on the AF-NS identified highly-common novel sequences shared among at least four, five and six samples**

**Supplementary Table 8. Annotation of the AF-NS identified novel sequence placements from long-read data. A list of the insertion sites, with placement locations, site type, IDs of samples containing the placement, whether in centromere, whether in telomere, intersecting genes, gene types, and repeat types of insertion regions**
